# BCMA/CD19 dual-targeting with FasTCAR: AZD0120 demonstrates potent preclinical efficacy in multiple myeloma

**DOI:** 10.64898/2026.06.05.730120

**Authors:** Huan Shi, Wenjie Yin, Hua Zhang, Xiaoyan Jiang, Jiaping He, Guangyao Zhu, Michael G. Overstreet, Mark Cobbold, Lianjun Shen

## Abstract

Chimeric antigen receptor (CAR)-T cell therapy has improved outcomes for patients with multiple myeloma (MM), but its broader use is restricted by manufacturing complexities and treatment-related toxicities. AZD0120 is a dual-targeting B-cell maturation antigen (BCMA)/CD19 CAR-T cell therapy manufactured via the rapid FasTCAR process. We developed a dual-targeting “loop” CAR that incorporates a novel humanized anti-BCMA single-chain variable fragment (scFv), clone SG, and an FMC63-derived anti-CD19 scFv. This AZD0120 CAR preserved functional binding to both antigens and conferred robust in vitro and in vivo cytotoxicity while maintaining single-antigen reactivity.

Conventional manufacture of CAR-T cells with the AZD0120 CAR (AZD0120C) yielded cells with minimal tonic signaling, limited responsiveness to soluble BCMA, and preservation of naïve/stem cell memory-enriched phenotypes, yet robust cytokine production upon BCMA^+^ target engagement. AZD0120C demonstrated cytotoxicity comparable to benchmark BCMA CAR-Ts across MM lines in vitro and showed strong in vivo expansion and tumor control in xenograft models.

FasTCAR manufacturing – designed to shorten vein-to-vein timelines and enrich less-differentiated phenotypes – further enhanced in vivo performance: AZD0120 consistently achieved superior tumor control and greater CAR-T expansion vs AZD0120C across disseminated MM.1S, NALM-6, and JeKo-1 models, with superior efficacy observed at lower cell doses. Collectively, these data support clinical evaluation of AZD0120 as a differentiated BCMA/CD19 CAR-T cell therapy with the potential to improve disease control and patient access in MM.

**Key Points:**

- AZD0120 is a dual-targeting CAR-T that displays a favorable anti-myeloma functional profile and co-targets a source of potential relapse
- The FasTCAR process yields T_N/SCM_-rich CAR-T populations, promotes in vivo expansion and achieves potent tumor control in xenograft models

## Introduction

Multiple myeloma (MM) is an incurable hematologic malignancy characterized by the proliferation of malignant plasma cells in the bone marrow. Despite significant advances in treatment, most patients eventually relapse, highlighting the critical need for therapeutic strategies that achieve deeper and more durable responses.^1,2^

Chimeric antigen receptor (CAR)-T cell therapy has emerged as a potentially transformative immunotherapy in MM, enabling targeted elimination of malignant plasma cells.^3,4^ B-cell maturation antigen (BCMA) is currently the most clinically validated target in MM as it is predominantly expressed on differentiated B cells, including malignant plasma cells. BCMA-directed CAR-T cell therapies were first approved by the U.S. Food and Drug Administration and European Medicines Agency in 2021, and have demonstrated high response rates with minimal residual disease-negativity in heavily pretreated patients with MM.^3,4^ While efficacy has been encouraging, some BCMA CAR-T products have been associated with serious adverse events, including non-immune effector cell-associated neurotoxicity syndrome (non-ICANS) neurologic complications that can be irreversible or even fatal.^5–9^ Furthermore, though some patients achieve durable responses for >1 year, most patients eventually relapse,^2^ potentially pointing to escape of a disease progenitor population. Thus, there remains a need for a differentiated therapy to further improve patient outcomes.

Patients with MM harbor circulating clonotypic CD19^+^ B cells that are genetically related to the malignant plasma cell clone.^10–16^ These clonotypic B cells carry chromosomal aberrations also found in the malignant plasma cell clone^17–20^ and may represent a disease-propagating population that contributes to relapse, particularly under therapeutic pressure.^19^ CD19 has been detected at low levels on malignant plasma cells^21^ and its expression is associated with treatment resistance and poor survival,^22^ supporting its rationale as a complementary target in MM. Importantly, a clinical study showed that antibody-mediated depletion of CD19-expressing cells in patients with MM reduced myeloma colony-forming units in the bone marrow.^23^ We and others have shown that combining BCMA and CD19 CAR-T cells led to a greater reduction in colony-forming units in bone marrow from patients with MM in vitro.^24,25^ These data have supported the design of clinical studies demonstrating the safety and efficacy of CD19 CAR-T administered with or without BCMA CAR-T in both relapsed/refractory MM and in high-risk newly diagnosed MM post-transplant.^24,26–28^ Together, these preclinical and clinical data support CD19 as a complementary target to BCMA and show that dual-targeting or co-administered BCMA/CD19 CAR-T can more consistently suppress myeloma progenitors and improve disease control across MM settings than targeting BCMA alone.

Translating these advances into consistent real-world benefit requires addressing the bottleneck of conventional CAR-T production, where extended lead times and capacity limits can impede prompt delivery. This delay increases the risk of interim disease progression and can potentially compromise patient eligibility. Next-generation manufacturing strategies, such as the FasTCAR platform, aim to overcome these limitations by significantly shortening CAR-T production timelines. The FasTCAR process uses a single “concurrent activation-transduction” step with lentiviral vectors, enabling efficient and stable CAR expression without the need for ex vivo expansion.^29,30^ The resulting CAR-T cells exhibit enhanced proliferative capacity and tumor killing activity and are enriched for less-differentiated phenotypes, including naïve and stem cell memory (T_N/SCM_) subsets.^29,30^ Importantly, enrichment for T_N/SCM_ populations has been associated with superior anti-tumor responses in preclinical studies^31–33^ and improved long-term clinical outcomes.^34–37^

AZD0120 is a BCMA/CD19 dual-targeting CAR-T cell therapy manufactured using the FasTCAR platform. Here, we describe the preclinical characterization of AZD0120 with the aim of validating a novel single-chain variable fragment (scFv) binder and dual-targeting loop CAR design, comparing its phenotype and function against clinical benchmarks, and demonstrating the advantages conferred by rapid FasTCAR manufacturing in the development of an effective and accessible CAR-T cell therapy for MM.

## Methods

### Cell lines and genetic engineering

MM.1S, RPMI8226, NALM-6, JeKo-1, JJN3, KMS12, KMS34, U226B1, NCIH929, SUDHL10, SUPB15, HeLa, K562, and HUH7 cell lines were obtained from American Type Culture Collection (Manassas, Virginia, USA), Japanese Collection of Research Bioresources (Ibaraki, Japan), and Leibniz Institute DSMZ-German Collection of Microorganisms and Cell Cultures GmbH (Braunschweig, Germany). Where indicated, tumor lines expressed luciferase for imaging. HeLa and K562 cells expressing BCMA, CD19, or both were generated by lentiviral transduction and sorted for high antigen expression. All cell lines were mycoplasma negative (MycoAlert, Mycoplasma Detection Kit, Lonza, Basel, Switzerland) and short tandem repeat authenticated.

### Surface plasmon resonance

Binding kinetics of the clone SG anti-BCMA scFv to recombinant human BCMA (ACRO Biosystems, Beijing, China) were measured on a Biacore 8K (GE Healthcare, Arlington Heights, Illinois, USA) with 1:1 Langmuir fitting to determine K_D_, k_a_, and k_d_.

### CAR constructs and lentiviral vectors

AZD0120 is a dual-target “loop” CAR structure comprising anti-CD19 (FMC63-derived) and clone SG, a novel high-affinity anti-BCMA scFv, arranged in a loop configuration upstream of a CD8α hinge/transmembrane domain, 4-1BBζ co-stimulatory domain, and CD3ζ. Single-target CAR designs used the same backbone. Some CARs used a CD28ζ costimulatory domain, where indicated. Self-inactivating lentiviral vectors (EF1α promoter) were produced by transient transfection of HEK293T cells, clarified (0.45 μm), concentrated by ultracentrifugation, and stored at −80°C. Functional titers were determined by flow-based transduction. Benchmark comparator BCMA CAR-T were reverse engineered from sequences obtained from the International Nonproprietary Names (INN) database (INN numbers 10906 [scFv CAR], 11131 [tanVHH CAR], and 12425 [DD CAR]) and cloned into lentiviral vectors.

### T cell isolation, manufacture, and culture

Peripheral blood mononuclear cells from healthy donor leukapheresis were isolated by Ficoll-Paque (GE Healthcare Life Sciences, Chicago, Illinois, USA) and T cells were enriched by negative selection (Pan T cell isolation kit; Miltenyi, Bergisch Gladbach, Germany). For conventional manufacture (AZD0120C and comparator CAR-T; AZD0120C denotes conventional manufacture of CAR-T cells with the AZD0120 CAR), isolated T cells were activated with with anti-CD3/CD28 Dynabeads (Thermo Fisher, Waltham, Massachusetts, USA); bead:cell 1:1, 48 hours) in X-VIVO 15 (Lonza) + 1% human serum albumin + interleukin (IL)-2 (300 IU/mL). Beads were then removed, cell density was adjusted to 10^6^ cells/mL, and cells were transduced with the lentiviral vector, and expanded for at least 9 days maintaining 0.5–1.5 × 10^6^ cells/mL.

For FasTCAR manufacture (AZD0120), a single-step concurrent activation–transduction process with lentiviral vectors was used per published methodology.^29^ Briefly, Dynabeads were added directly to isolated T cells for 30 minutes, followed immediately by addition of the lentiviral vector. The following day, beads were removed and cells were washed and cryopreserved. For assessment of CAR expression, transduced cells were maintained in X-VIVO 15 with 1% human serum albumin + IL-2 (300 IU/mL) for 4 days, and stained for fluorescence-activated cell sorting.

### Flow cytometry and CAR detection

Cells were stained with LIVE/DEAD^TM^ Fixable Aqua Dead Cell Stain Kit (Thermo Fisher) and labeled in phosphate buffered saline + 2% bovine serum albumin + 3.6 mM EDTA. Antibodies included CD45, CD3 (UCHT1), CD4 (SK3), CD8 (SK1), CD45RA (HI100), CCR7 (3D12), CD25, CD27, CD62L (DREG-56), CD137, PD-1 (EH12.2H7), TIM-3 (F38-2E2), LAG-3 (7H2C65), and TIGIT (A15153G). CAR detection used biotin-CD19 (ACRO Biosystems) with streptavidin-APC, BCMA-PE (ACRO Biosystems), or anti-FMC63 scFv (BioSwan Laboratories, Shanghai, China) at 4°C. Data were acquired on BD LSRFortessa/BD FACSymphony/BD FACSCanto II (BD Biosciences, Franklin Lakes, New Jersey, USA) and analyzed with FlowJo v10 (BD Biosciences) or Cytobank (Beckman Coulter, Brea, California, USA).

### In vitro cytotoxicity and cytokine analysis

Luciferase-expressing target cells were co-cultured with CAR-T cells at the indicated effector-to-target (E:T) ratios for 6–72 hours in 96-well plates. Following co-culture, plates were centrifuged and 50 µL of supernatant was collected for cytokine analysis. The remaining co-culture was resuspended, and 100 µL was transferred to a white 96-well plate, followed by the addition of 100 µL of luciferase substrate (ONE-Glo™ Luciferase Assay System, Promega, Madison, Wisconsin, USA). Luminescence was read on a SpectraMax i3x (Molecular Devices, San Jose, California, USA) and percent killing calculated as 100 × (target only relative luminescence unit [RLU]-sample RLU)/target only RLU (triplicates).

For real-time impedance assays, HeLa, HeLa-CD19, HeLa-BCMA, or HeLa-CD19-BCMA were seeded at 10^4^ cells/well on E-Plate 96 (xCELLigence RTCA MP [Agilent, Santa Clara, California, USA]). CAR-T cells were added the next day; impedance was recorded every 15 minutes and analyzed per manufacturer’s instructions. For degranulation and intracellular cytokine staining, CAR-T cells and targets were plated at E:T 1:2 with anti-CD107a, GolgiStop, and GolgiPlug (BD Biosciences) for 4 hours. Cells were stained for CD3 and CD8 as described, then analyzed by flow cytometry.

For repetitive stimulation assays, CAR-T cells were co-cultured with irradiated K562, K562-CD19, K562-BCMA, or K562-CD19/BCMA (20–40 Gy) cells every 3 days; viable CAR^+^ T-cell numbers and target clearance were quantified at each restimulation.

For stimulation with soluble BCMA (sBCMA), CAR-T cells were co-cultured with BCMA-null HUH7 target cells ± recombinant BCMA-His (ACRO Biosystems, 890 ng/mL) or with engineered BCMA-expressing HUH7 cells; cytokines were measured after 24 hours. Supernatants were analyzed for IL-2, IL-4, IL-6, IL-10, TNF-α, and/or IFN-γ using cytometric bead array (BD Biosciences) or electrochemiluminescence (Meso Scale Discovery, Rockville, Maryland, USA).

### Cell avidity (z-Movi)

Target monolayers were formed on poly-L-lysine–coated acoustofluidic chips (LUMICKS, Amsterdam, Netherlands). CAR-T cells labeled with CellTracker Deep Red C34565 (Thermo Fisher) were incubated for 15 minutes; acoustic force ramps were applied and detachment recorded. Dissociation force metrics were computed with z-Movi software (LUMICKS).

### In vivo studies

Female NOG (NOD.Cg-*Prkdc^scid^IL2rg^tm1Sug^*/JicCrl) or NOG-dKO (NOD.Cg-*Prkdc^scid^IL2rg^tm1Sug^B2m^em1Tac^H2-Ab1^tm1Doi^*/JicCrl) mice (6–10 weeks old; Vital River, Beijing, China) were used for animal studies conducted under the Institutional Animal Care and Use Committee-approved protocols.^38^ Mice were inoculated with tumor cells as indicated and then received CAR-T cells IV. In the MM.1S model, mice received 10^7^ tumor cells IV and were dosed with CAR-T cells 6–10 days later. In the RPMI8226 model, mice were implanted with 10^7^ tumor cells subcutaneously (SC) and dosed with CAR-T cells 17–18 days later. In the dual-tumor model, mice were first implanted with RPMI8226 (10^7^ cells SC), followed 14 days later by MM.1S (10^7^ cells IV); CAR-T cells were administered 4 days later, and complete responders were subsequently rechallenged with RPMI8226 (10^7^, SC) on study day 55. In the NALM-6 model, mice received 10^6^ tumor cells IV and CAR-T cells 6 days later. In the JeKo-1 IV model, mice received 3 × 10^6^ tumor cells IV and CAR-T cells 7 days later. For IV models, tumor burden was monitored twice weekly by bioluminescence imaging using an in vivo imaging system (IVIS), either IVIS Lumina III or IVIS Spectrum BL (PerkinElmer, Shelton, Connecticut, USA), per the manufacturer’s instructions. In SC models, tumor volumes were measured by calipers twice weekly, and blood collection and flow cytometry were performed once weekly. Baseline tumor burden (total flux by IVIS or tumor volume in mm³ by caliper) was assessed one day before or on the day of CAR-T dosing (defined as Day 0). Tumor-bearing animals were randomized based on baseline tumor burden, either manually or using Provantis (Instem, Stone, United Kingdom).

### Jurkat reporter assay

Jurkat triple reporter cells that express cyan, green, and red fluorescent proteins under control of NF-kB-, NFAT-, and AP-1-responsive elements, respectively, were transduced with CAR constructs and assayed for basal reporter activity without BCMA stimulation.

### Statistical analysis

Statistical analysis was carried out using GraphPad Prism Software version 10 (GraphPad Software, Boston, MA). Two-group comparisons used unpaired two-tailed t-tests. Multiple groups were analyzed by one-way or two-way analysis of variance with Tukey/Šidák correction. *P* < .05 was considered statistically significant.

## Results

### Characterization and functional activity of the novel BCMA-binding scFv CAR

We identified an antibody clone specific for human BCMA from a rat hybridoma library and humanized the clone to create clone SG, for which no off-target binding was identified (Figure S1). Surface plasmon resonance binding kinetics demonstrated that clone SG exhibited high affinity for human BCMA (K_D_: 6.78e-10 M, K_a_: 1.98e6 1/Ms, K_d_: 1.34e-3 1/s; Figure 1A). When incorporated as an scFv into a CAR construct, clone SG CAR-T cells demonstrated robust cytotoxic function against engineered HeLa cells expressing BCMA (Figure 1B). Furthermore, clone SG CAR-T cells degranulated and secreted cytokines in response to BCMA^+^ tumor cells, comparable to or exceeding the levels secreted from a benchmark scFv CAR comparator (Figure 1C–D). Importantly, clone SG CAR-T cells induced effective lysis of multiple tumor cell lines expressing BCMA, demonstrating anti-tumor activity similar to the scFv comparator CAR (Figure 1E).

**Figure 1.**
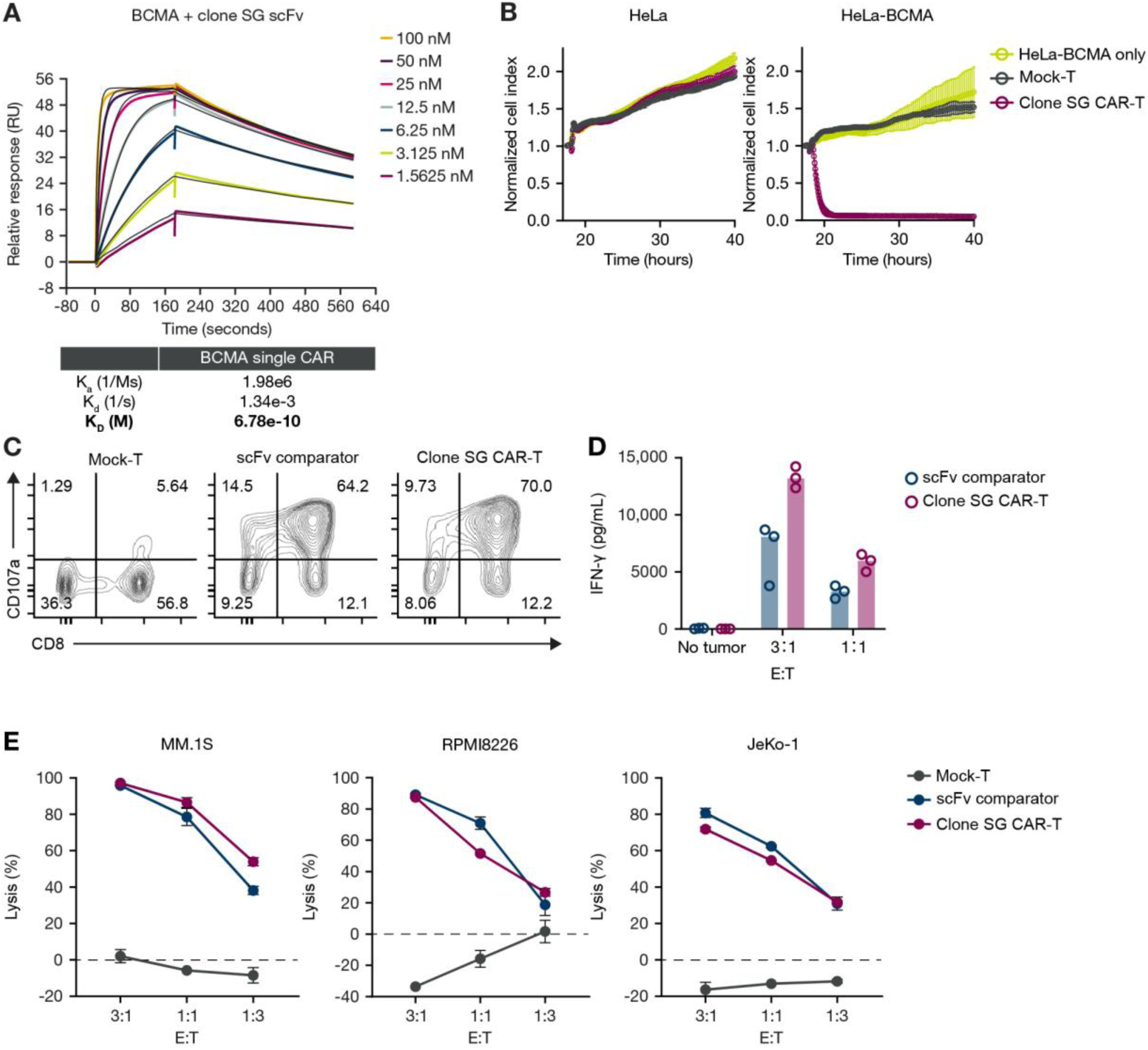
Description and activity of anti-BCMA clone SG scFv/CAR. (A) SPR binding kinetics of clone SG BCMA binder to human BCMA. (B) Cytotoxic function of clone SG in CAR format (CD28ζ costimulatory domain) against engineered HeLa cells expressing BCMA. (C, D) Degranulation and cytokine secretion of clone SG CAR-T cells (4-1BBζ costimulatory domain) compared to scFv CAR comparator after co-culture with MM.1S cells. (E) Cytolysis of BCMA^+^ tumor cell lines by clone SG CAR-T cells compared to the scFv CAR comparator. RU, Response Unit; SPR, surface plasmon resonance.

### In vitro and in vivo activity of the AZD0120 CAR

The AZD0120 CAR was designed to target both BCMA^+^ and CD19^+^ and incorporates binding domains for each target antigen within a single loop configuration (Figure 2A). In vitro studies showed that the T cells expressing the AZD0120 loop CAR killed target cells faster than CAR-T with the same scFvs in a tandem CAR format (Figure S2A). Functional binding to both target antigens was validated with Jurkat cells expressing the AZD0120 CAR binding recombinant BCMA and CD19 (Figure 2B), and simultaneous binding of both antigens was confirmed by surface plasmon resonance (Figure S2B). Importantly, affinity to BCMA was maintained in the loop CAR structure (K_D_: 5.41e-10 M, K_a_:2.48e6 1/Ms, K_d_: 1.34e-3 1/s). CAR-T cells expressing the AZD0120 CAR using a conventional manufacturing process (AZD0120C; Figure 2C) exhibited higher avidity than the single CAR-T controls when encountering dual-antigen–positive targets (BCMA and CD19), suggesting potentially stronger antigen recognition (Figure S1C). Against single-antigen–positive cell lines, cellular avidity was comparable to that of the single CAR control (Figure S2C).

**Figure 2.**
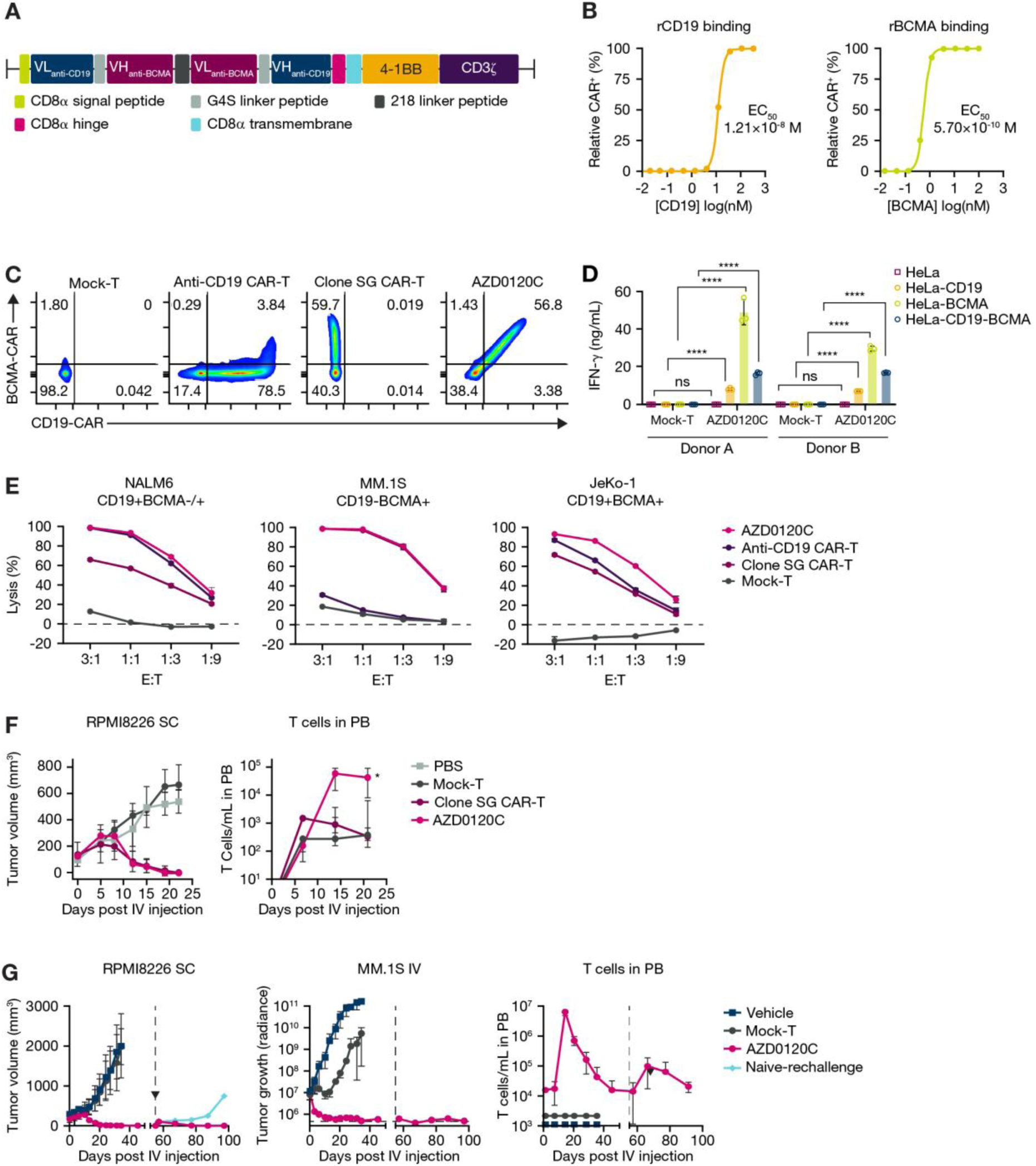
Description and in vitro and in vivo activity of AZD0120 CAR. (A) Gene structure of BCMA^+^CD19 AZD0120 loop CAR. (B) Binding of recombinant CD19 or BCMA to Jurkat cells expressing AZD0120C CAR. (C) Staining patterns of CAR-T expressing single- or dual-BCMA and/or CD19 CARs. (D) Cytokine production by CAR-T expressing AZD0120 CAR or single CARs after co-culture with engineered HEK target cells expressing BCMA and/or CD19. (E) In vitro cytotoxicity against BCMA^+^ and/or CD19^+^ cell lines at 6 hours. (F) Anti-tumor activity and in vivo T cell expansion of clone SG CAR-T cells (CD28ζ costimulatory domain) vs AZD0120C in subcutaneous model of RPMI-8226 (NOG mice; Day -17: 10^7^ RPMI-8226 SC, Day 0: 3.5×10^6^ CAR-T IV). (G) Anti-tumor activity and in vivo T cell expansion of AZD0120C in dual tumor challenge model (subcutaneous RPMI-8226 and disseminated MM.1S) with subcutaneous rechallenge with RPMI-8226 (NOG-dKO mice; Day -18: 10^7^ RPMI-8226 SC, Day -4: 10^6^ MM.1S IV, Day 0: 5e6 CAR-T IV, Day 55: rechallenge with 10^7^ RPMI-8226 SC). *, *P <* .05 by ANOVA Holm-Šidák test of AZD0120C vs clone SG; ****, *P <* .0001 by two-way ANOVA; HEK, human embryonic kidney; ns, not significant; PB, peripheral blood; PBS, phosphate buffered saline; r, recombinant; VH, variable heavy chain; VL, variable light chain.

Functionally, AZD0120C CAR-T cells produced pro-inflammatory cytokines upon co-culture with engineered HEK target cells expressing BCMA and/or CD19, indicating effective activation in response to either antigen and demonstrating preserved dual-target functionality compared with single CAR controls (Figure 2D). AZD0120C similarly demonstrated anti-tumor activity against BCMA^+^ and CD19^+^ cell lines in vitro (Figure 2E). In a subcutaneous xenograft model of RPMI-8226, AZD0120C demonstrated anti-tumor activity comparable to clone SG CAR-T cells but AZD0120C cells expanded 64-fold more than clone SG cells at day 14 (Figure 2F). To assess its efficacy in a more complex disease setting, we utilized a dual tumor model comprising subcutaneous RPMI-8226 and disseminated MM.1S. Building on single model results, AZD0120C demonstrated potent anti-tumor activity coupled with strong T cell expansion, with sustained protection after a subcutaneous rechallenge with RPMI-8226 (Figure 2G).

### Phenotypic and functional in vitro profile of AZD0120C against clinical benchmarks

We conducted an in vitro comparison of AZD0120C against leading clinical benchmark constructs. AZD0120C preserved the T_N/SCM_ compartment at levels similar to non-transduced T cells and more so than tanVHH and DD comparators (Figure 3A–B). In cytotoxicity assays against a panel of luciferase-tagged tumor cell lines, AZD0120C demonstrated similar cytotoxic potency to the benchmark CAR-T constructs across different time points (24, 48, and 72 hours) and E:T ratios (only E:T 1 shown for simplicity in Figure 3C).

**Figure 3.**
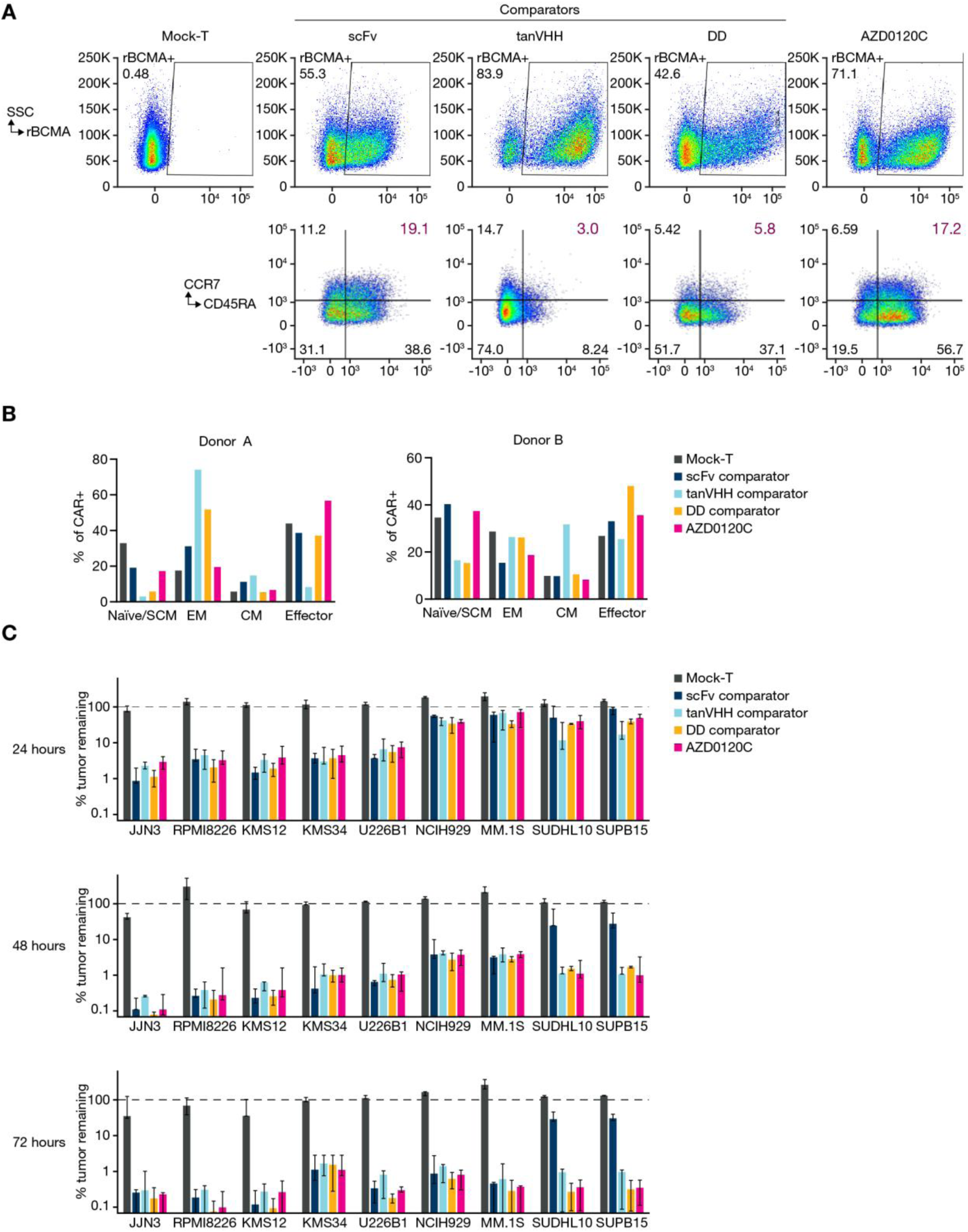
In vitro memory subsets and anti-tumor activity against comparators. (A) FACS analysis of AZD0120C vs comparators shown from a single donor. CAR expression shown by binding recombinant BCMA (first line) and memory subsets (second line). (B) Summary of T cell memory compartment analysis in Panel A for two donors. Naïve/T_SCM_: CD45RA^+^CCR7^+^, Effector memory: CD45RA^−^CCR7^−^, Central memory: CD45RA^−^CCR7^+^, Effector: CD45RA^+^CCR7^−^. (C) CAR-T cells were co-cultured with the indicated luciferase-tagged tumor cell lines at an E:T 1 and harvested at the indicated time points. Data are presented as % luciferase signal relative to tumor cells without T cells. Data shown are medians +/− range for 3 unique donors (DD comparator 2 donors only). CM, central memory; EM, effector memory; FACS, fluorescence-activated cell sorting.

Phenotypic analysis of CAR-T cells in culture (no co-culture with tumor cells) showed that AZD0120C CAR-T cells displayed low levels of CD25 and CD137 relative to comparator CAR constructs, indicating minimal tonic signaling (Figure 4A–B). Consistent with these data, Jurkat reporter cells expressing AZD0120C showed minimal reporter activation in the absence of antigen relative to comparator CAR constructs, further indicating an absence of antigen-independent CAR signaling (Figure 4C; additional data in Figure S3). Consistent with the differences observed in surface phenotype, AZD0120C released minimal IFN-γ in the absence of BCMA^+^ target cells or in the presence of soluble BCMA (sBCMA), unlike benchmark CAR-T constructs that released greater amounts of IFN-γ under both conditions (Figure 4D). Importantly, AZD0120C produced high levels of IFN-γ in response to BCMA^+^ target cells, demonstrating robust anti-tumor potential, and the highest levels of IL-2, consistent with enrichment of T_N/SCM_ phenotype cells (Figure 4E).

**Figure 4.**
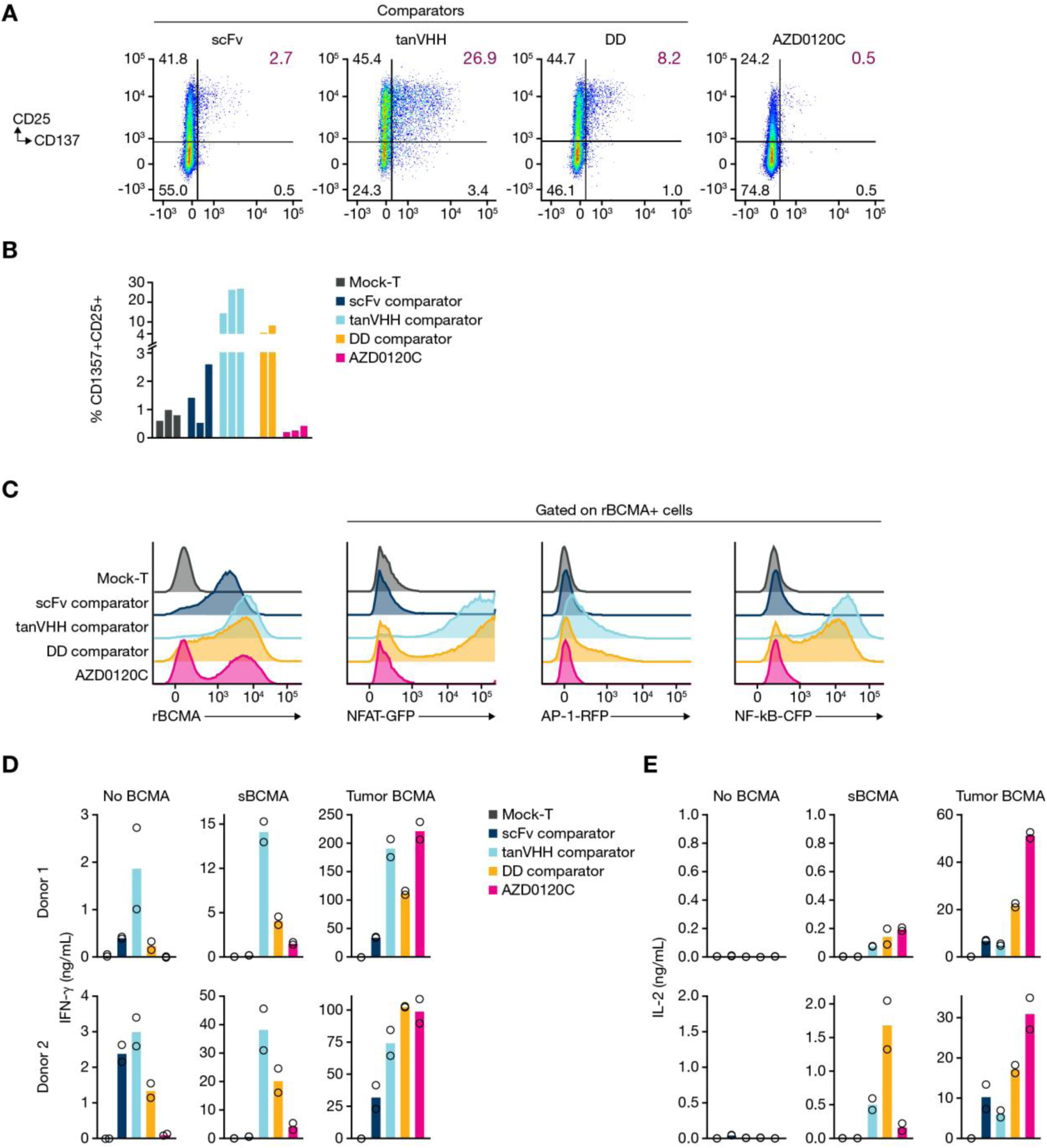
Tonic signaling and response to sBCMA. (A) FACS analysis of CD25 vs CD137 for AZD0120C vs comparators as indicators of tonic signaling. (B) Summary of CD137^+^CD25^+^ analysis in Panel A for three unique donors (DD comparator utilized for two donors only). (C) Jurkat reporter cells transduced to express the indicated CARs were stained and analyzed for reporter expression in the absence of BCMA (no co-culture). (D) CAR-T cells were cultured with BCMA(−) HUH7 tumor cells, HUH7 tumor cells plus sBCMA (150 nM), or engineered BCMA^+^ HUH7 tumor cells. Culture supernatants were then assayed for IFN-γ production after 24 h. Data shown for two independent donors. Bars represent the mean of the experimental replicates. (E) IL-2 production readout for experiment in Panel D. CFP, cyan fluorescent protein; GFP, green fluorescent protein; RFP, red fluorescent protein.

### In vitro characteristics of AZD0120 cells

The FasTCAR process is designed to both preserve a favorable T_N/SCM_ phenotype and shorten manufacturing times. Because of the rapid stimulation, transduction, and cryopreservation, CAR expression cannot be immediately detected. We therefore assessed CAR expression kinetics and early readouts post-manufacture. Analysis of AZD0120 FasTCAR cells from three donors 4 days post-activation and CAR transduction showed that CAR^+^ cells were preferentially enriched for T_N/SCM_ subsets compared with the bulk of the T-cell population, indicating preservation of a less-differentiated T-cell state early after manufacture (Figure 5A). Due to this inherent lag in CAR expression associated with the FasTCAR process, AZD0120 cells were not immediately cytotoxic in early tumor-killing assays (24 h) and only showed clear killing after 48- and 72-h assays (Figure 5B). However, once stimulated in vitro with K562 cells expressing BCMA and/or CD19, AZD0120 cells expanded more robustly than AZD0120C cells (Figure 5C) and demonstrated similar cytolytic function in cytotoxicity assays against BCMA^+^ and/or CD19^+^ cell lines (Figure 5D).

**Figure 5.**
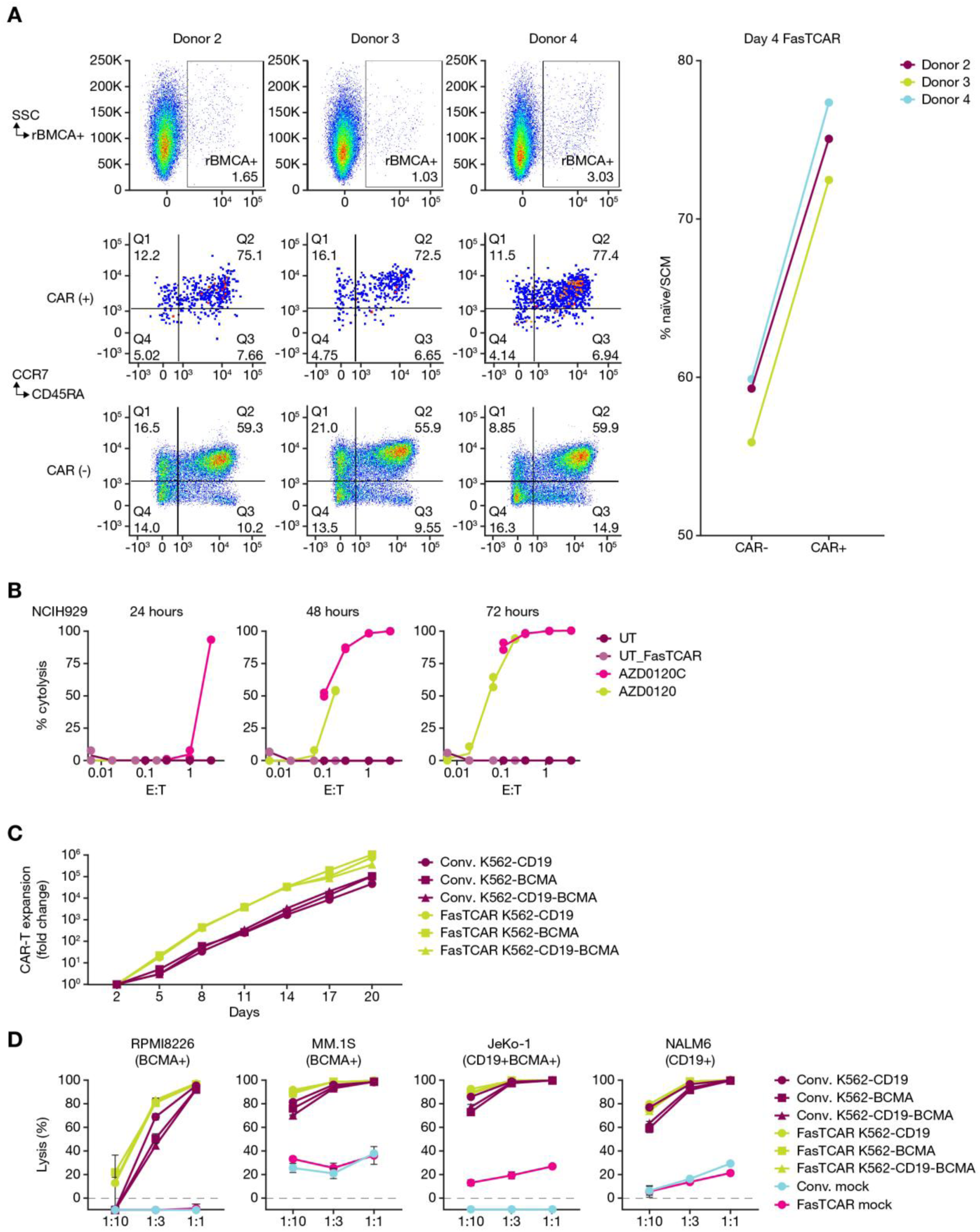
FasTCAR cells in vitro. (A) FACS analysis of AZD0120 FasTCAR cells from three donors 4 days after activation and CAR transduction. CAR expression (rBCMA binding, top line), memory subset analysis on CAR^+^ cells (middle line) and CAR-cells (bottom line). Graph on right shows that CAR^+^ cells at Day 4 are preferentially T_N/SCM_ compared to bulk T-cell population. (B) In vitro cytotoxicity assay against NCI-H929 comparing AZD0120C and AZD0120 (FasTCAR) from a single donor, harvested at three different time points. E:T ratios were set using CAR staining on day of co-culture for AZD0120C. For AZD0120 (0% CAR^+^), 50% CAR% was used for assay setup, then back-calculated using Day 4 staining to determine adjusted E:T. (C) AZD0120C or AZD0120 were co-cultured with K562 cells expressing BCMA, CD19, or both every 3 days. Relative CAR-T cell expansion is shown. (D) AZD0120C or AZD0120 were co-cultured with K562 cells as in panel C for 6 days. CAR-T cells were then co-cultured with the indicated cell lines and assayed for cytotoxicity. Conv., conventional.

### In vivo activity of AZD0120 FasTCAR cells

To determine the impact of FasTCAR manufacturing on anti-tumor efficacy, the in vivo efficacy of AZD0120 was directly compared to AZD0120C across various disseminated tumor models at low cell doses that were suboptimal for AZD0120C. In a disseminated MM.1S model, AZD0120 provided more robust and durable tumor control across a range of doses compared to AZD0120C. This enhanced anti-tumor activity was consistently associated with significantly greater T cell expansion in the blood of treated mice (Figure 6A; Figure S4). Similar superior efficacy and T cell expansion for AZD0120 were observed in disseminated NALM-6 (Figure 6B; Figure S5) and JeKo-1 (Figure 6C; Figure S6) models.

**Figure 6.**
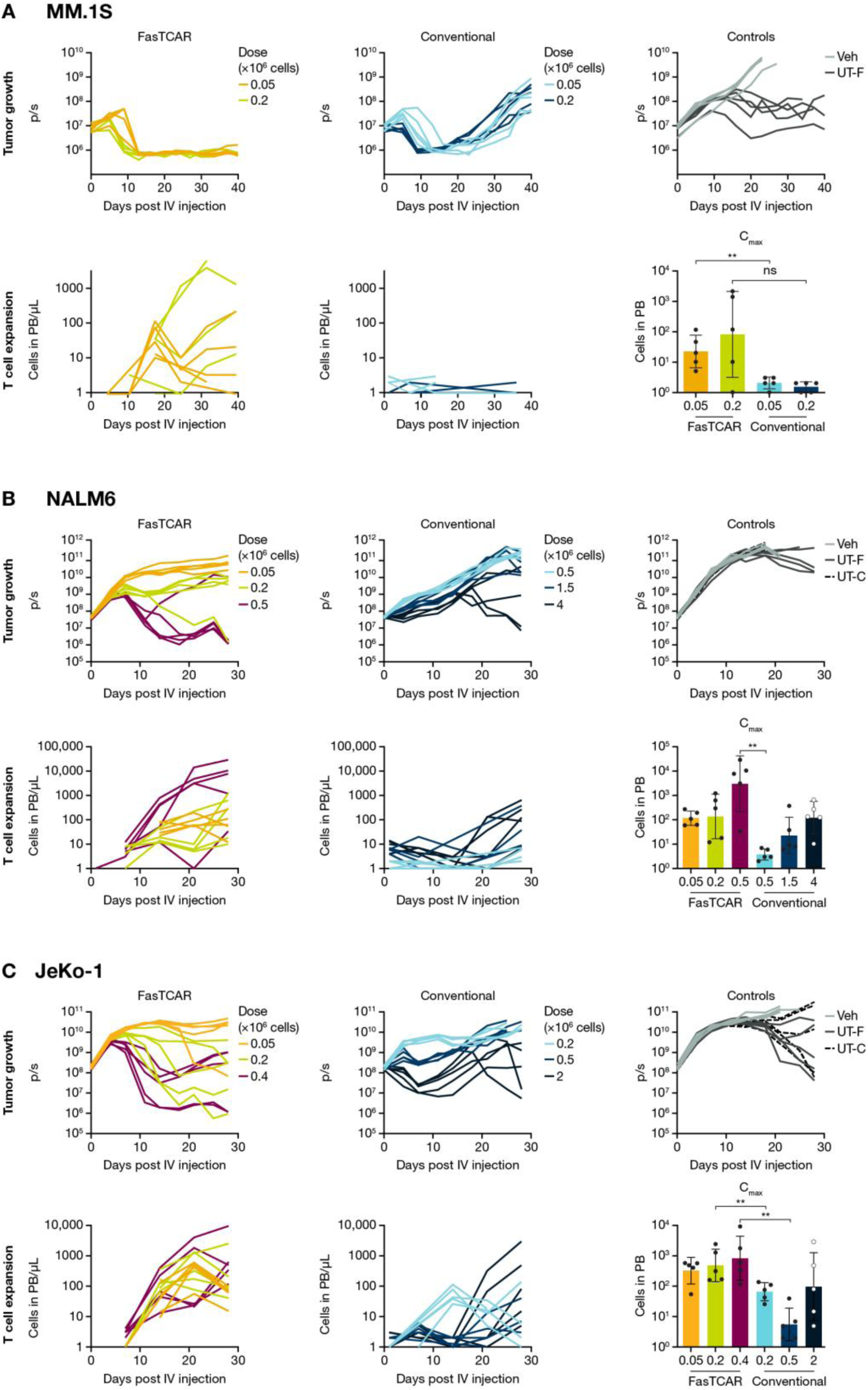
Profile of FasTCAR vs conventional cells *in vivo.* (A) In vivo disseminated MM.1S model comparing AZD0120C and AZD0120 at indicated doses. Luciferase signal shown represents tumor growth. Graphs on right shown T cell expansion in blood from individual mice, colors matched to graphs on left (Day -10: 10×10^6^ MM.1S IV, Day 0: 0.05 or 0.2×10^6^ CART-T [for both AZD0120C and AZD0120] IV). (B) In vivo disseminated NALM-6 model comparing AZD0120C and AZD0120 at indicated doses. Layout and graphs same as in panel A (Day -6: 1×10^6^ NALM-6 IV, Day 0: 0.05, 0.2, and 0.5×10^6^ CAR-T for AZD0120 and 0.5, 1.5, and 4×10^6^ for CAR-T for AZD0120C, all IV). (C) In vivo disseminated JeKo-1 model comparing AZD0120C and AZD0120 at indicated doses. Layout and graphs same as in panel A (Day -7: 3×10^6^ JeKo-1 IV, Day 0: 0.05, 0.2, and 0.4×10^6^ CAR-T for AZD0120 and 0.2, 0.5, and 2×10^6^ for CAR-T for AZD0120C, all IV). **, *P <* .01; ns, not significant by Mann-Whitney test comparing FasTCAR and conventional CAR-T at equivalent dose levels; Veh, vehicle.

## Discussion

Our preclinical studies describe a novel anti-BCMA scFv binder that, when incorporated into a novel dual-targeting loop CAR structure with an anti-CD19 scFv, exhibits potent antimyeloma activity while maintaining a favorable functional profile. In particular, the AZD0120 CAR showed high selectivity for BCMA without detectable off-target binding, in contrast to reports of off-target interactions associated with other benchmark CAR constructs.^39^ This selectivity may contribute to a more controlled activation profile and is hypothesized to reduce unintended effects.

AZD0120C was characterized by a low level of tonic signaling relative to benchmark CAR-T constructs. This, combined with a lack of detectable off-target binding, may result in a more favorable therapeutic profile. Excessive tonic signaling is associated with constitutive activation and cytokine secretion, which may contribute to increased inflammatory risk.^40^ This is particularly relevant, as late-onset neurotoxicity has been associated with high peak CAR-T expansion, cytokine release syndrome (CRS)/ICANS, and high-level CAR-T persistence with some BCMA-directed CAR-T cell therapies.^6,41^

Consistent with this controlled activation profile, AZD0120C demonstrated minimal release of IFN-γ when exposed to sBCMA despite robust cytokine production in the presence of BCMA^+^ tumor cells. This finding contrasts with benchmark CAR-T products that released cytokines in the absence of BCMA or in response to sBCMA. Lack of activation of AZD0120 by sBCMA may reduce the risk of cytokine-mediated toxicities such as CRS and off-tumor inflammation. Recent data highlight the potential for antigen-induced CAR multimerization of a benchmark CAR-T, including cross-linking via soluble antigen to amplify CAR signaling,^42^ underscoring the relevance of this observation. Together, these data indicate that AZD0120C couples favorable on-tumor effector function with tighter control of off-tumor and antigen-independent activation.

Phenotypically, AZD0120C preserved a substantial T_N/SCM_ phenotype compared to benchmark constructs. This less-differentiated state, observed independently of the FasTCAR process, suggests that the intrinsic design of the AZD0120 CAR contributes to maintaining a younger and more potent T-cell population, potentially related to its minimal tonic signaling. Enrichment of T_N/SCM_ subsets has been associated with improved proliferative potential, sustained anti-tumor responses, and favorable clinical outcomes across CAR-T platforms.^32,34–37^ While our data support favorable phenotypes, formal linkage to long-term persistence and clinical durability will require prospective correlative analyses.

AZD0120 cells show a delayed cytolytic response to BCMA^+^ tumor cells, attributable to the gradual CAR expression following lentiviral vector transduction. This delayed surface CAR expression provides a conceptual framework for more predictable timing and lower intensity of CRS that can be evaluated clinically. A gradual reduction in tumor burden during in vivo expansion may blunt a systemic cytokine response at peak expansion compared with conventionally manufactured CAR-T cells, when a bolus dose of highly-activated effector CAR-T cells encounters a larger tumor burden, possibly leading to a more synchronous and intense release of cytokine. AZD0120 also shows an enrichment for a T_N/SCM_ phenotype, which is associated with a less inflammatory activation profile.^32^

A unique strength of AZD0120 lies in its dual targeting of both BCMA and CD19. While our data confirm comparable or greater efficacy against BCMA^+^ MM cells relative to benchmark constructs, the targeting of CD19^+^ progenitor B cells is hypothesized to provide greater durability by eliminating a potential source of relapse or antigen escape. Although the specific long-term durability benefit of eliminating CD19^+^ cells cannot be directly demonstrated in the current study, the hypothesis is well supported in the literature^21,23,24,43^ and will be explored in ongoing clinical trials.

The FasTCAR platform shortens manufacturing timelines and enriches for less-differentiated T cell phenotypes. This rapid CAR-T cell manufacturing process can substantially shorten the interval between patient diagnosis and CAR-T cell infusion. Beyond logistical advantages, the results here show that AZD0120 FasTCAR cells exhibit superior expansion and more durable tumor control compared with conventionally manufactured CAR-T cells across multiple disseminated tumor models. By shortening production timelines and enriching for less-differentiated T-cell states, the FasTCAR process may also help improve treatment accessibility and therapeutic control. While these findings support the hypothesis that AZD0120 could reduce off-tumor activation and potentially mitigate cytokine-driven toxicities, formal safety and durability studies will require prospective clinical evaluation, including assessment of cytokine kinetics and the contribution of CD19 targeting to relapse prevention. Ongoing studies are designed to test these hypotheses and to define the clinical impact of AZD0120 in patients with MM.

## Acknowledgments

We thank Jiajing Wen, Jessica Tong, Dominique Brown, and Quincy Barton for technical expertise and execution of the in vitro studies. We thank Tao Wang and Suying Zhang for technical expertise and execution of the in vivo studies.

This work was supported by AstraZeneca.

Medical writing support under the direction of the authors, was provided by Angharad Morgan, PhD, CMPP, an employee from the Publications and Medical Affairs Division of Omnicom Health Medical Communications, and was funded by AstraZeneca, in accordance with Good Publication Practice (GPP 2022) guidelines.

## Authorship

### Contribution

H.S., W.Y., H.Z., and L.S. conceptualized the study; H.S., W.Y., and H.Z. developed the methodology; H.S. and W.Y. curated the data, performed the formal analysis, and provided project administration, study resources, and validation; H.S., G.Z., and H.Z. carried out the investigation; H.S., J.H., W.Y., M.G.O., and L.S. supervised the research; H.S., J.H., W.Y., and M.G.O. visualized the data, and wrote the original draft of the article; and G.Z., M.C., H.S., J.H., W.Y., H.Z., M.G.O., and L.S. wrote and reviewed subsequent drafts of the article.

### Author contributions

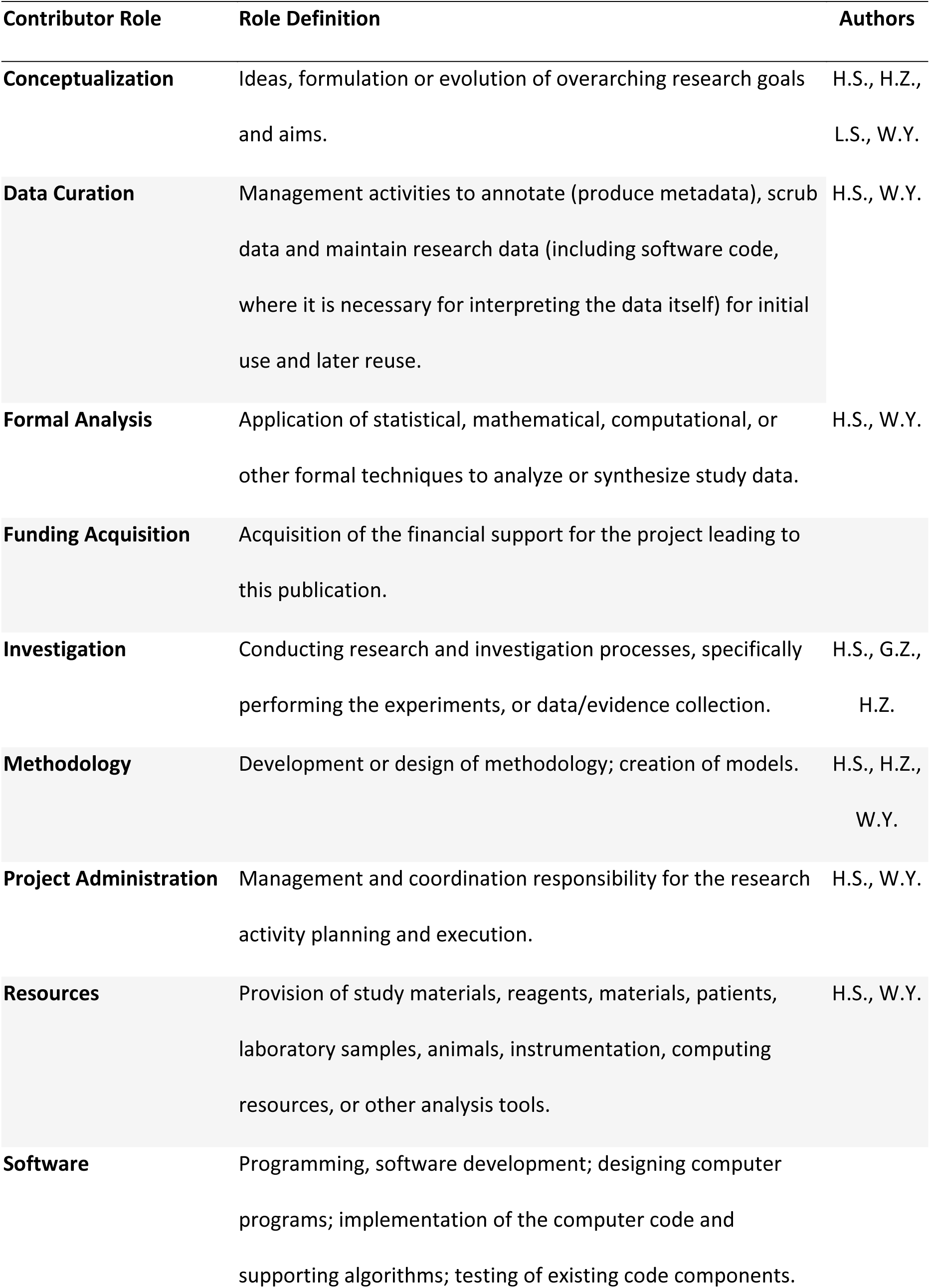
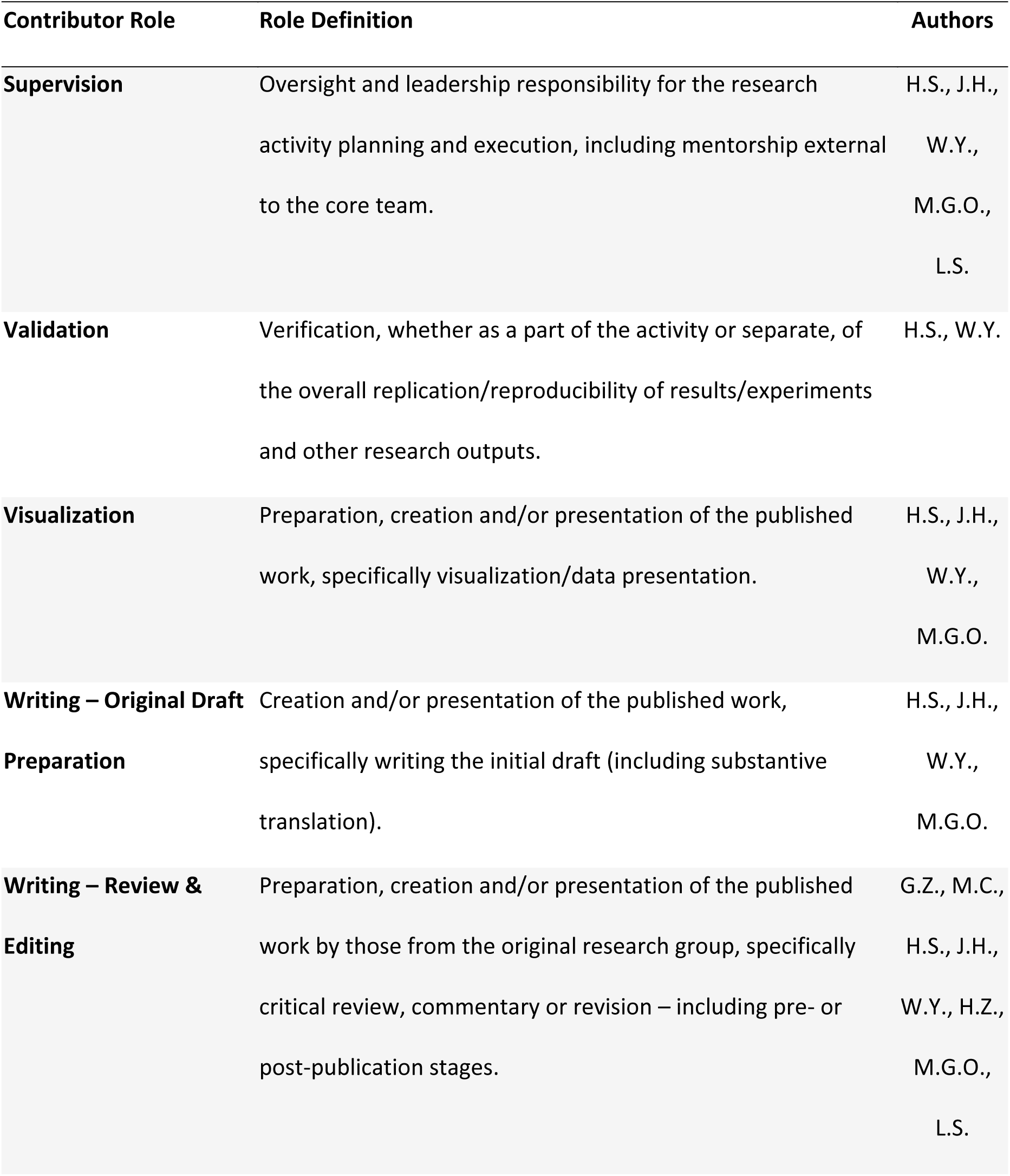

---

**Figure S1.**
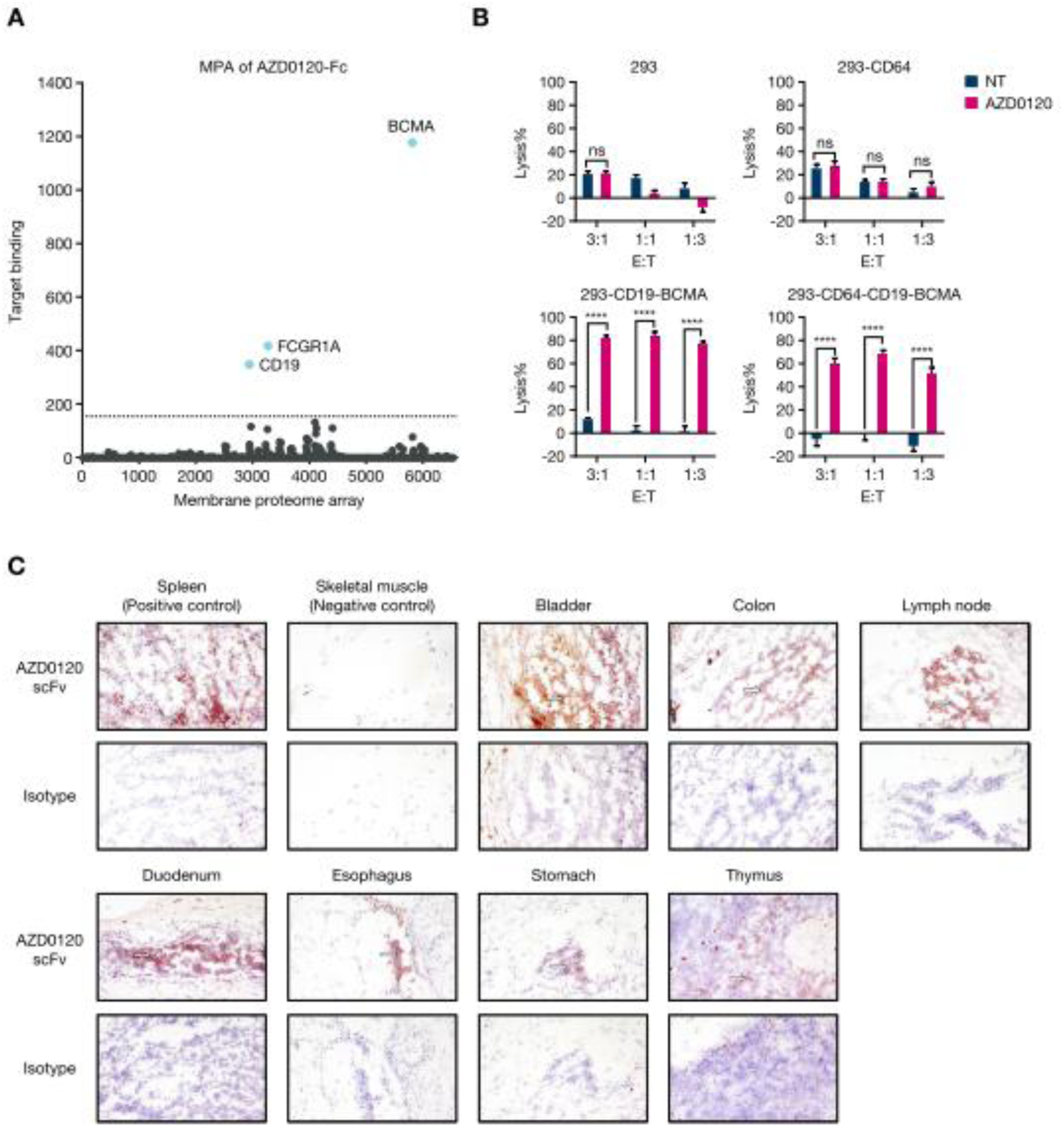
Off target evaluation of AZD0120 CAR. (A) Membrane Proteome Array (MPA) results of AZD0120-Fc. (B) Lysis of target cells by AZD0120 or its matching NT-F at different E:T for 20 h. Percentage of lysis was expressed as changes in Cell Index relative to the Cell Index of the target only at one specific time point. Tests were repeated in triplicate. ****, *P <* .0001 by two-way ANOVA. (C) Cross-reactivity of AZD0120-Fc with human frozen tissues. Each tissue tested was from three individual donors. Arrows indicate positive membrane staining. CAR, chimeric antigen receptor; E:T effector to target ratio; ns, not significant; NT-F, N-terminal fragment.

**Figure S2.**
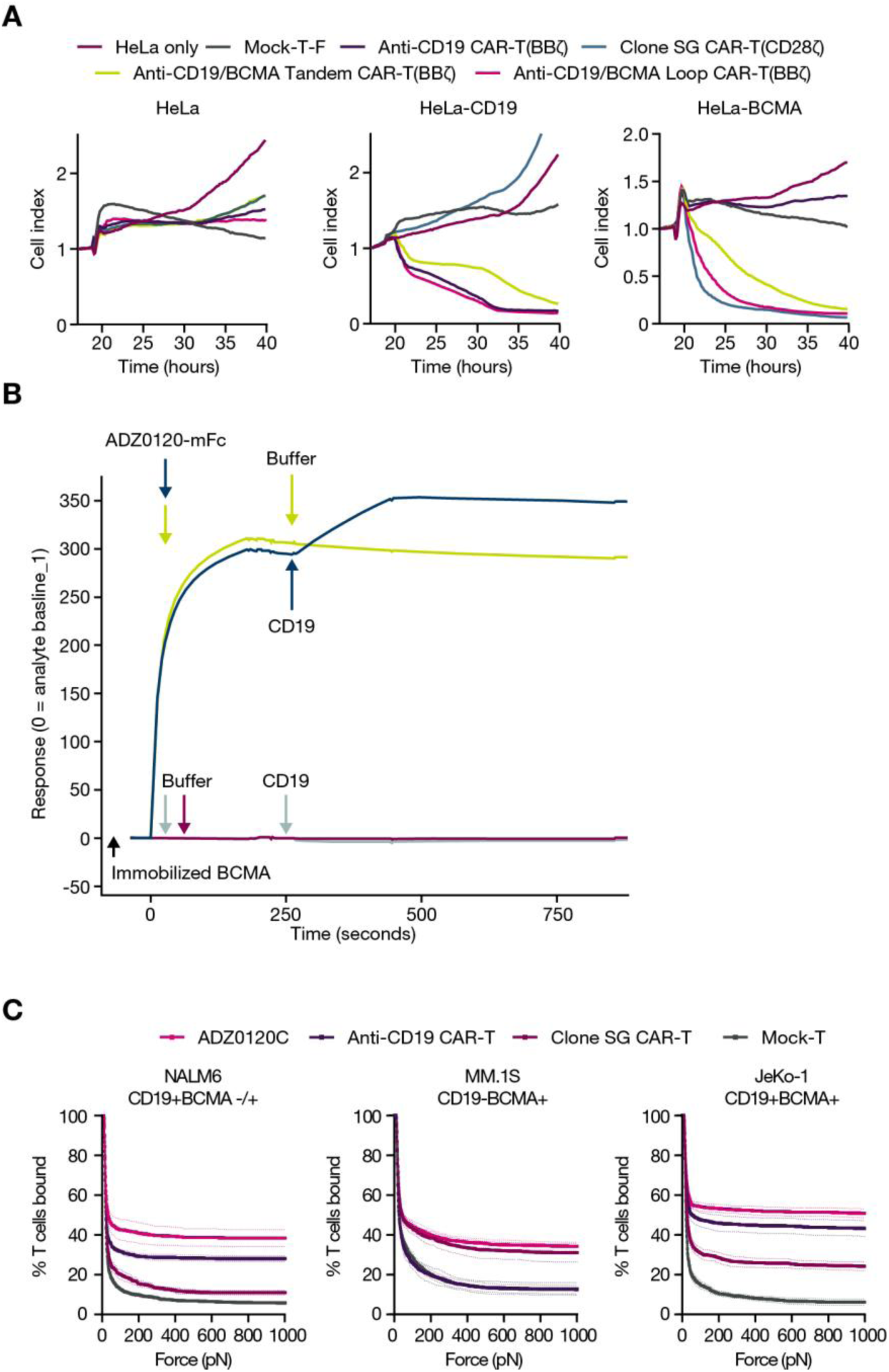
Activity of the Loop BCMA-CD19 CAR. (A) In vitro cytotoxicity assays of single- or dual-CAR-T cells against engineered HeLa cells expression BCMA or CD19. (B) Surface plasmon resonance kinetic binding summary of AZD0120 CAR to immobilized BCMA and then soluble CD19. (C) Lumicks cellular avidity data for ADZ0120 or single CAR-T binding to the indicated tumor cell lines. BCMA, B-cell maturation antigen; CAR, chimeric antigen receptor.

**Figure S3.**
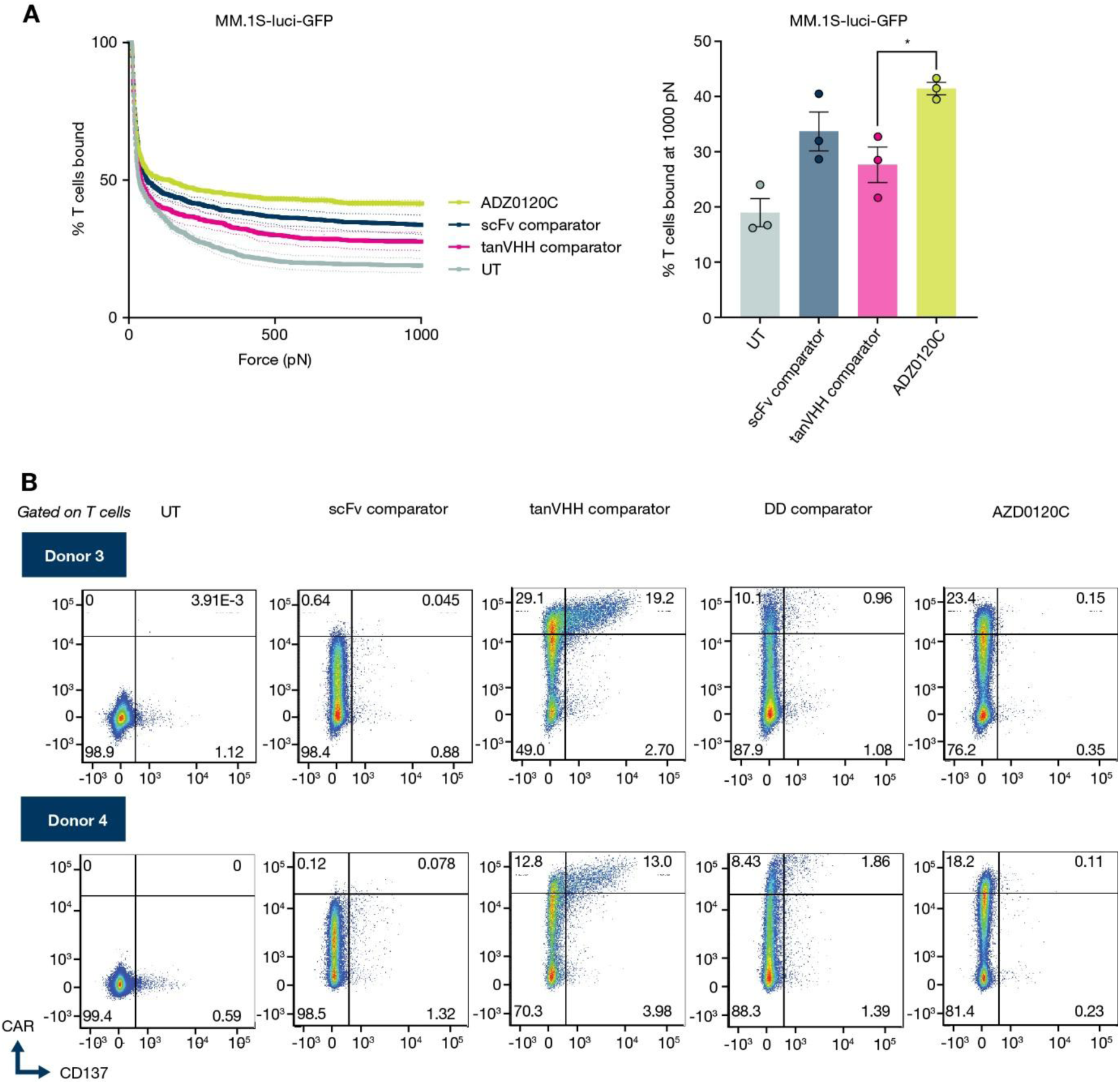
Cellular avidity and tonic signaling. (A) Lumicks cellular avidity data for ADZ0120C CAR-T on MM.1S tumor cells vs comparators. (B) Expression of CD137 on CAR^+^ cells from AZD0120C and comparators. Quadrant gate set to highlight restricted expression of CD137 to cells expressing highest levels of CAR. CAR, chimeric antigen receptor; GFP, green fluorescent protein; Q, quadrant; rBCMA, recombinant B-cell maturation antigen; scFv, single-chain variable fragment.

**Figure S4.**
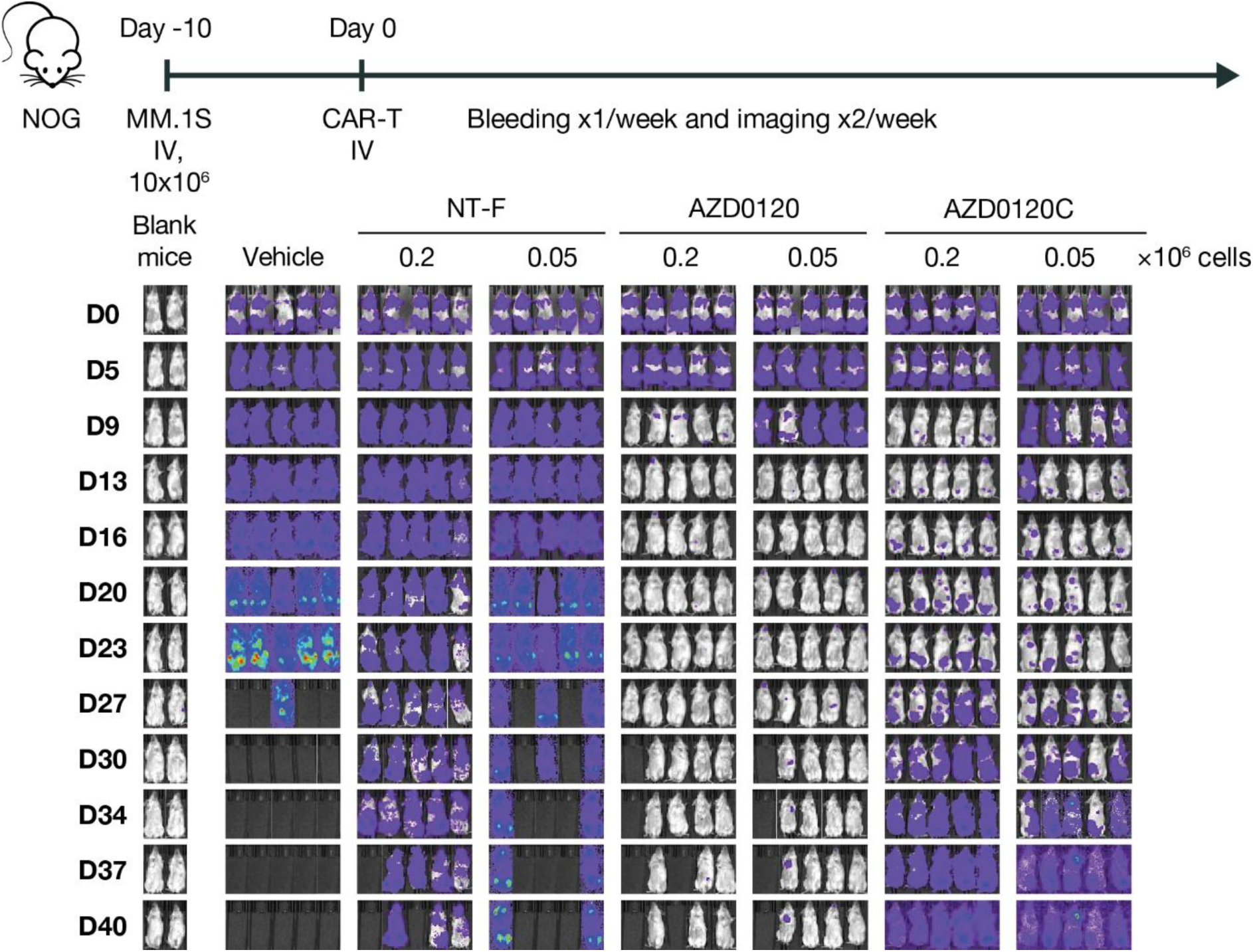
Schematic of disseminated MM.1S tumor model and IVIS image time course comparing control non-transduced FasTCAR T cells (NT-F), AZD0120, and AZD0120C CAR-T (corresponding to Figure 6A). CAR, chimeric antigen receptor; D, Day; IVIS, in vivo imaging system.

**Figure S5.**
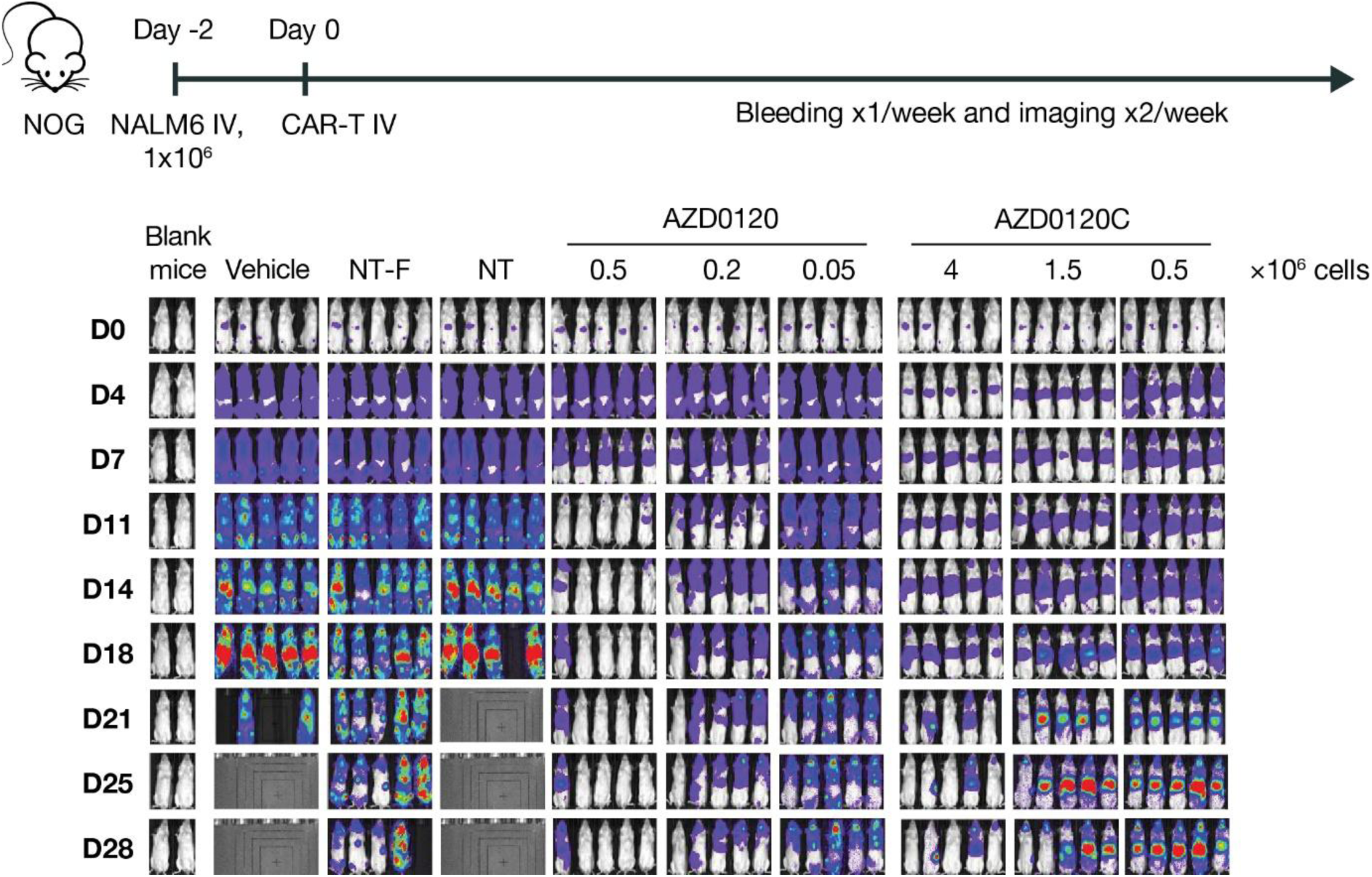
Schematic of disseminated NALM-6 tumor model and IVIS image time course comparing control non-transduced FasTCAR T cells (NT-F), non-transduced T cells (NT), AZD0120, and AZD0120C CAR-T (corresponding to Figure 6B). CAR, chimeric antigen receptor; D, Day; IVIS, in vivo imaging system.

**Figure S6.**
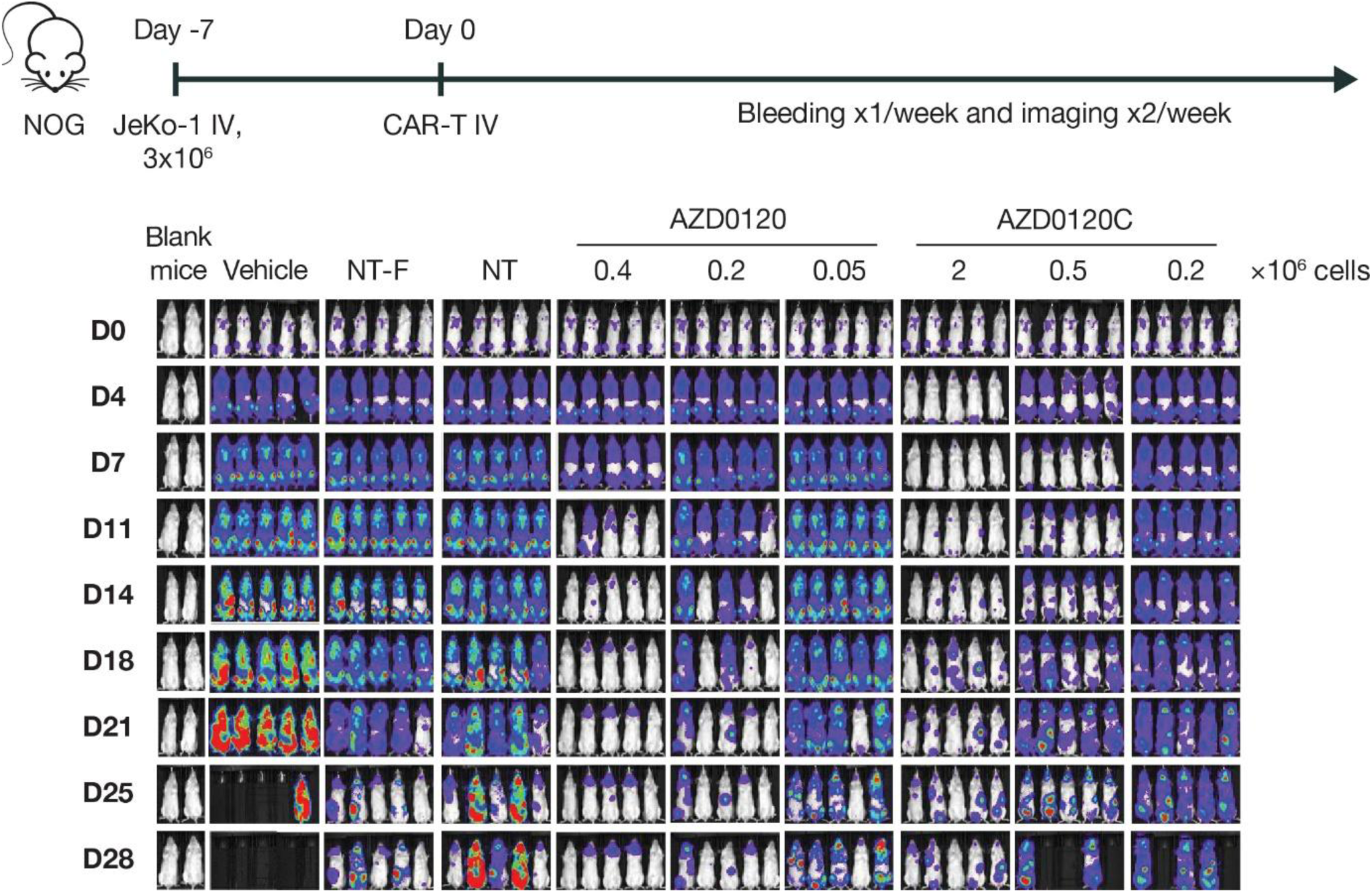
Schematic of disseminated JeKo-1 tumor model and IVIS image time course comparing control non-transduced FasTCAR T cells (NT-F), non-transduced T cells (NT), AZD0120, and AZD0120C CAR-T (corresponding to Figure 6C). CAR, chimeric antigen receptor; D, Day; IVIS, in vivo imaging system.

